# Berberine chloride ameliorated PI3K/Akt-p/SIRT-1/PTEN signaling pathway in insulin resistance syndrome-induced rats

**DOI:** 10.1101/459610

**Authors:** Marwa El-Zeftawy, Doaa Ghareeb, Rasha Saad, Salma Mahmoud, Nihal Elguindy, Mohammed El-Sayed

## Abstract

Insulin resistance is one of dangerous factors as it leads to numerous metabolic disorders such as non-insulin dependent diabetes mellitus. It affects most tissues mainly adipose tissue, liver and muscle. Nowadays, berberine has several medical applications against diseases. The current study was carried out to identify the effect of berberine chloride (BER-chloride) on phosphatidyl inositol-3-kinase/ phosphorylated protein kinase B/ sirtuin type 1/ phosphatase and tension homologue (PI3K/Akt-p/SIRT-1/PTEN) pathway during insulin resistance phenomena. Insulin resistance model was performed in experimental rats by using high fat diet. Plasma glucose, serum insulin, lipid profiles, hepatic oxidative stress markers were estimated. Serum transaminases activities and kidney function tests were determined. Further, hepatic PI3K, AKt-p, SIRT-1; PTEN levels were assayed. The concentration of adiponectin in serum, hepatic tissue and white adipose tissue was determined. Moreover, fold change in hepatic insulin, insulin receptor and retinol binding protein-4 (RBP4) at molecular level was performed. Histopathological study of white adipose tissue was also determined. The results showed increase the rats’ body weights, blood glucose, homeostatic model assessment, glycated hemoglobin, insulin and lipid profiles levels in group of rats fed on high fat diet for eight weeks and this elevation was decreased after administration of BER-chloride for two weeks. Further, BER-chloride administration exhibited improvement of oxidative stress parameters, PI3K, AKt-p, SIRT-1 and PTEN. This was associated with down-regulation of RBP4. According to these data we conclude that, BER-chloride mediated several insulin signaling pathways that could be of therapeutic significance to insulin resistance.

## 1. Introduction

The increased prevalence of obesity between individuals has become a serious health problem worldwide. Under normal conditions, β-cell of pancreas maintains the normal glucose tolerance by increasing insulin release to overcome the reduction of insulin efficiency. One of the predisposing risk factor to obesity is the amount of fat in the diet due to modern life styles. Obesity usually accompanied by insulin resistance and hyperglycemia [1]. Insulin resistance defined as a disease condition in which insulin is secreted from β-cell of pancreas but its function is impaired in peripheral tissues such as liver, adipose tissue and skeletal muscle. Insulin resistance usually associated with metabolic disorders such as hyperlipidemia, type II diabetes mellitus, non-alcoholic fatty liver and cardiovascular disease and early mortality is considered one of insulin resistance prognosis in some individuals [2]. Phosphatidyl inositol-3-kinase/ phosphorylated protein kinase B (PI3K/Akt) pathway is one of the most important signaling pathways which involved in metabolic effect of insulin [3].

Therefore any treatment strategy of insulin resistance should be associated with targeting of insulin signaling pathway complications. Nowadays, using of herbal compounds occupied a huge importance in medical field.

Berberine (BER) is a natural isoquinoline alkaloid isolated from different plants such as *Berberis vulgaris* [4]. BER is a strong base which is usually unstable when present in free form so it usually accompanied with chloride ion in form of BER-chloride [5, 6]. BER has several pharmacological activities, it acts as anticancer [6, 7], antiinflammatory [7, 8], antileishmanial [8, 9] and anti-human immunodeficiency virus [5, 10]. BER can be used for improving some cardiac diseases and intestinal infections especially bacterial diarrhea [9, 10]. Furthermore, recently it is used as a neuroprotective agent against some neurodegenerative diseases such as *Alzhimer’s* and *Parkinson’s* diseases as it has the ability to pass the blood brain barrier [4].

To investigate whether BER-chloride has a protective effect on insulin resistance, we set up *in vivo* model for insulin resistance by High fat diet (HFD) feeding. The effects of BER-chloride on various insulin signaling pathway were investigated.

## 2. Materials and methods

### 1.1. Materials

BER-chloride was obtained from Sigma-Aldrich Chemical Co. (USA). Kits and reagents for the assay of blood glucose level (BGL), protein, lipid profiles [total cholesterol (TC), triacylglycerol (TG) and high density lipoprotein-cholesterol (HDL-c)] and glycated hemoglobin (HbA1C), as well as both kidney function tests (creatinine and urea) and liver enzymes [alanine aminotransferase (ALT) and sspartate aminotransferase (AST)] were obtained from Spinreact (Spain), Human (Germany), Biosystem (Egypt), and Biolabo (France), respectively. Ribouncleic acid (RNA) extraction kit, Maxime reverse transcription (RT) premix kit, 2x Taq master mix, deoxyribonucleic acid (DNA) Ladder, ribonuclease (RNase)-free water, and the primer sequences of β-actin, insulin, insulin receptor (IR) and rat retinol binding protein-4 (RBP4) were obtained from Qiagen (Germany), Intron Biotechnology (Korea) and Fermentas, Thermo fisher scientific (Germany), respectively.

Enzyme linked immunoassay (ELISA) kit of insulin, PI3K, AKt-P, sirtuin type 1 (SIRT-1), phosphatase and tension homologue (PTEN) and adiponectin were purchased from DRG (USA), Wuhan Fine Biological Technology Co. (China), Ray Biotech (Georgia), MyBiosource (USA), Abcam (USA), Bioscience (USA), respectively. Foline reagent, thiobarbituric acid (TBA), reduced glutathione (GSH), 5,5′-dithio-bis-2-nitrobenzoic acid (DTNB), cumene hydroxide, methyle green, sodium pentobarbital and poly ethylene glycol (PEG) were obtained from Sigma-Aldrich Chemical Co. (St. Louis, Mo, USA). Organic solvents; ethanol 95% and methanol were of high pressure liquid chromatography (HPLC)-grade and brought from Merck (USA). Other reagents were obtained with high grade.

### 1.2. Experimental animal protocol and samples preparation

Female albino Sprague-Dawley rats (*Rattus norvegicus),* of body weight (130 – 150) g and aged (10 – 12) weeks old, were obtained from the experimental animal house of Medical Research Institute, Alexandria University, Egypt. Rats were housed in polycarbonate cages in groups of six rats per cage. They were kept under conventional conditions of temperature and humidity with a 12-h photoperiod. Food and water were supplied *ad libitum*. The experimental animals were conducted in accordance with the National Institutes of Health Guide for the Care and Use of Laboratory Animals (NIH 1996). This study was carried out in strict accordance with the recommendations in the Guide for the Care and Use of Laboratory Animals. The protocol was approved according to the Ethics of Animal House in Medical Technology Center, Alexandria University, Egypt.

All the animals were acclimatized for one week before the start of the experiment. After that, thirty animals were divided into five groups (n = 6 per group), Group 1 (Sham control group) that were healthy and free from any disease, rats of this group were fed standard diet for 10 weeks, Group 2 (Control vehicle) fed low fat diet (LFD) for eight weeks then received 20% PEG by intragastric tube (10 ml/kg Bwt) for two weeks [11, 12]. Group 3 (Control BER-chloride) fed LFD for eight weeks then orally given BER-chloride dissolved in 20% PEG (100 mg/kg Bwt) for two weeks. Group 4 (Induction group) fed HFD for eight weeks then orally given 20% PEG (10 ml/kg Bwt) for two weeks [11, 12]. Group 5 (Induction treated group) fed HFD for eight weeks then orally given BER-chloride dissolved in 20% PEG (100 mg/kg Bwt) for two weeks [13].

Rat’s body weights were recorded at the first and last week of treatment. Blood sampling and animal scarification were performed under sodium pentobarbital anesthesia, and all efforts were made to minimize suffering. At the end of the study, animals were fasted overnight after 8-h, blood samples were collected in sodium fluoride tubes for assessment of fasting BGL. After full fasting period (12-h), blood samples were collected, and then centrifuged at 3000 rpm for 10 min. The obtained serum was kept at −20°C until analyzed. Hepatic tissues and white adipose tissue from control and experimently groups were exised immediately and washed with ice-cold saline. Homogenization was carried out in 0.1M sodium phosphate-buffer, pH 7.4 (for hepatic tissue) and in 0.15M potassium chloride (for white adipose tissue). The homogenate was centrifuged at 4000 rpm for 15 min at 4°C and supernatant was stored at −80°C until analysis [14, 15]. In each group, part of liver was preserved in liquid nitrogen, and stored at −80°C for total RNA isolation and polymerase chain reaction (PCR) analysis and part of white adipose tissue was fixed in 10% neutral buffered formalin solution for histopathological examination.

### 1.3. Biochemical, molecular, histopathological studies and statistical analysis

#### 1.3.1. Biochemical and molecular studies

Glucose levels in all groups were measured by a glucose assay kit that is dependent on glucose oxidase-peroxidase method. Serum insulin of all groups was assayed by DRG insulin ELISA kit. The insulin resistance was evaluated by calculating the homeostatic model assessment-insulin ressistance (HOMA-IR) as previously described [16]. HbA1C % was determined by Biosystem kit. Total protein concentration was determined spectrophotometrically using Beirut assay based kit. Serum lipid profile (TC, TG, and HDL-c), kidney function tests (urea and creatinine) and liver function tests (ALT and AST) were carried out according to commercial kits manufacturer’s instructions. LDL-c and very low density lipoprotein cholesterol (vLDL-c) levels were calculated by using a specific formula [17]. Standardized methods were used to determine the level of thiobarbituric acid reactive substances (TBARS) [18] and GSH [19] in liver. In addition, hepatic activities of xanthine oxidase (XO) [20], glutathione peroxidase (GPx) [21, 22] and adenine triphosphatase (ATPase) [23] were carried out.

PI3K, Akt-P, SIRT-1 and PTEN levels in liver homogenate and adiponectin level in serum, liver homogenate and white adipose tissue were determined by using ELISA kits. These assays employ the quantitative sandwich enzyme immunoassay technique. Primers used for PCR technique were designed using the known sequences for the respective genes (Table 1). Programs are given as denaturation temperature/ denaturation times/ annealing temperature/ annealing times/ extension temperature/ extension times/ number of cycles. The primers were run on Mini Cycler (Eppendorf, Labcaire, Germany).

**Table 1.**
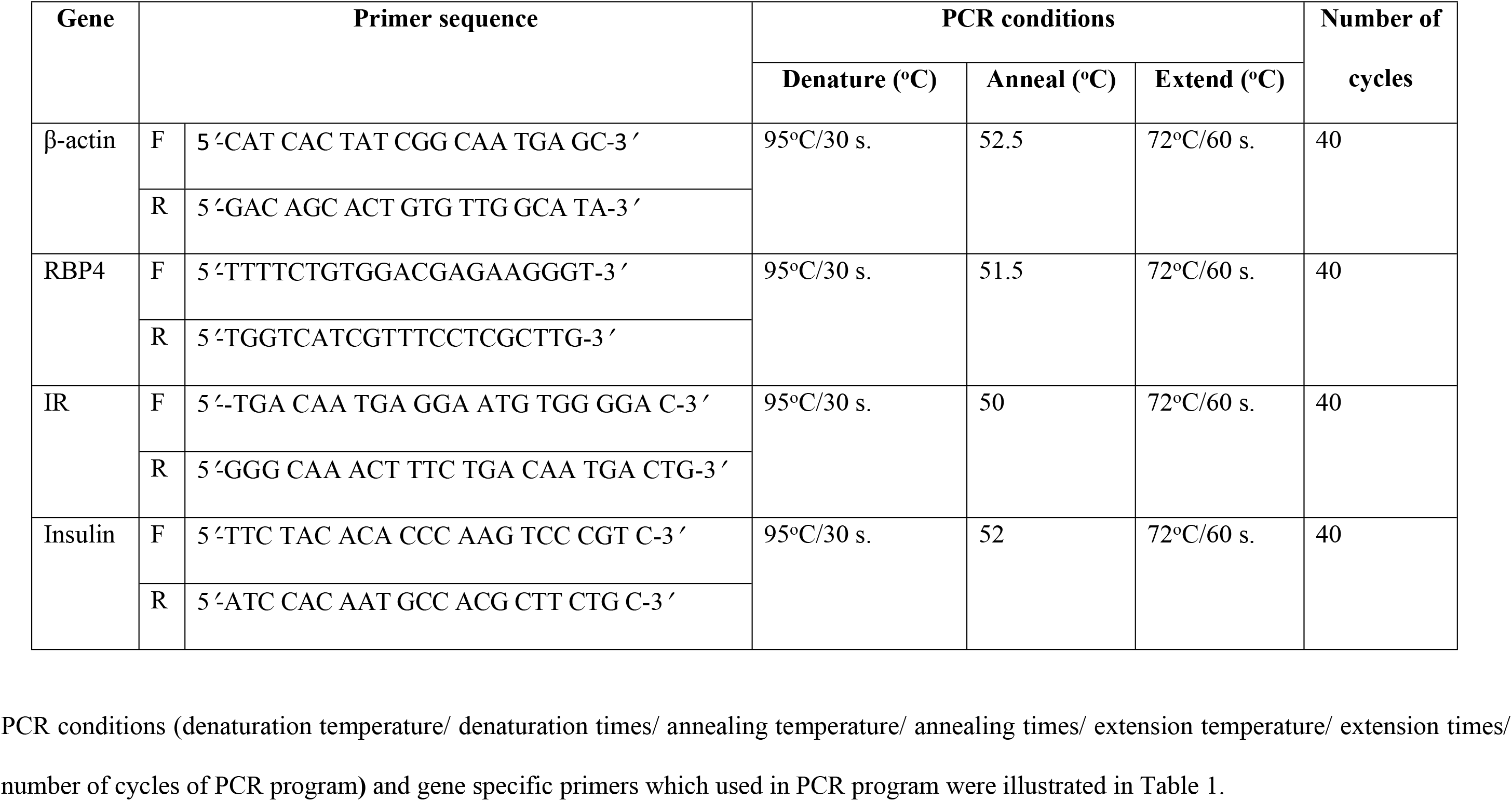
Primer sequences and PCR conditions

Total RNA was extracted from hepatic tissue by using total RNA extraction Kit and processed according to kit manufacturer’s instructions. After that the concentration of total RNA was measured by spectrophotometer at 260 and 280 nm. One microgram of the isolated RNA was reverse transcribed into single-strand complementary DNA (cDNA) using reverse transcriptase (Maxime RT Pre-Mix kit, Fermentas, EU). For gene expression, the gene specific primers were used and the programs (Table 1) were optimized for each primer pair and all programs started with a 30s period at 95°C and ended with a 60s extension at 72°C. The PCR products were resolved on 1.5% agarose gel. Gels were stained with ethidium bromide, visualized by 30 nm Ultraviolet Radiator (Alpha-Chem. Imager, USA), and photographic record was made. The optical density and the microgram content of bands were calculated by the UVIBAND MAX software program.

#### 2.3.2. Histopathological preparation of white adipose tissue

White adipose tissue of each rat from each group was excised and immediately fixed at 10% neutral buffered formalin solution after washing with ice cold normal saline. The resultant fixed tissue samples were used for histological examination in the Histopathology Laboratory of Medical Technology Center, Alexandria University, using the routine procedures developed in the respective laboratories. The tissue was cut at 3 mm thick, and the blocks were embedded in paraffin. Using a rotary microtome, sections of 8 μm thickness were cut. The sections were stained with hematoxylin and eosin and examined under Olympus microscope (Olympus, Tokyo, Japan) at (40X) magnification for any histopathological changes.

#### 2.2.3. Statistical analysis

Data were analyzed using Primer of Biostatistics software program (Version 5.0) by one-way analysis of variance (ANOVA). Significance of means ± SD was detected groups by using multiple comparisons Student-Newman-keuls test at p < 0.05. Adiponectin correlation was analyzed by SPSS (Version 20.0) software program, using person coefficient (r).

## 3. Results

### 3.1. Body weight, BGL, insulin resistance, lipid profiles, oxidative stress markers, serum transaminases activity and kidney function tests

#### 3.1.1. Body weight, BGL and insulin resistance parameters

Feeding of HFD for 8-weeks leads to increase the body weight of the rats than control level. Fasting BGL, HOMA-IR and HbA1C were also elevated 1.4, 3.7 and 45.2-folds, compared to sham control rats. Moreover, elevation of insulin level was reported in HFD group 1.0-fold in serum and 72.3-fold in hepatic tissue. Administration of BER-chloride leads to decrease fasting BGL, serum insulin, HOMA-IR and HbA1C to 0.5, 0.2, 0.6 and 0.2-folds, respectively compared with HFD rats (Table 2). Moreover, HFD up regulated insulin gene expression in liver tissue and the treatment with BER-chloride for two weeks did not showed any positive effect on insulin expression (Fig 1).

**Fig 1.**
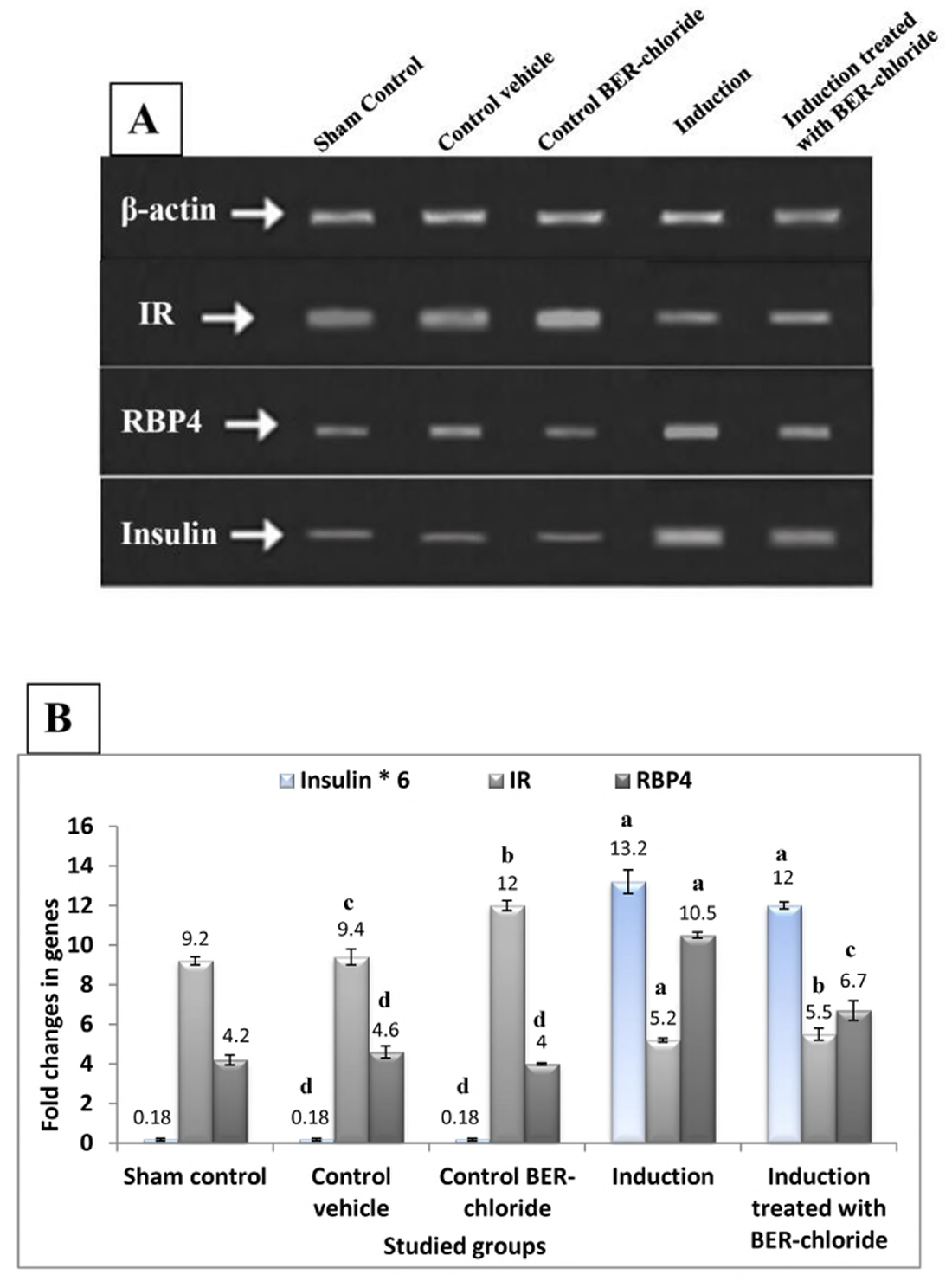
Effect of BER-chloride on the fold change of insulin, IR and RBP4 genes. (A) Agrose gel electrophoresis of gene expression of insulin (293 bp), IR (129 bp) and RBP4 (392 bp) compared to β-actin (300 bp). (B) Fold change of gene expression in liver homogenate after the treatment of diabetic induced experimental animals represented as 6 rats ± SE. ANOVA (one way) followed by Student-Newman-keuls test. Means with letters (a), (b), (c) and (d) were statistically represented compared to sham control group as a at p < 0.001, b at p < 0.01, c at p < 0.05 and d at p > 0.05.

**Table 2.** Effect of BER-chloride on body weight, BGL, serum insulin, HOMA-IR and HbA1C after the treatment of diabetic induced experimental animals.

|  | **Body weight (g)** | **BGL (mg/dl)** | **Serum insulin**<br>**(μIU/ml)** | **HOMA-IR (mg/dl)** | **HbA1C (%)** |
| --- | --- | --- | --- | --- | --- |
| **Sham control** | 141.0 ± 7.4<sup>a</sup> | 76.0 ± 1.4<sup>a</sup> | 10.0 ± 0.2<sup>a</sup> | 1.91 ± 0.061<sup>a</sup> | 4.2 ± 0.4<sup>a</sup> |
| **Control vehicle** | 142.2 ± 8.04<sup>a</sup> | 73.33 ± 2.6<sup>a</sup> | 10.1 ± 0.6<sup>a</sup> | 1.9 ± 0.13<sup>a</sup> | 4.4 ± 0.42<sup>a</sup> |
| **Control BER-chloride** | 144.0 ± 6.07<sup>a</sup> | 75.2 ± 3.31<sup>a</sup> | 10.45 ± 0.33<sup>a</sup> | 1.92 ± 0.12<sup>a</sup> | 4.0 ± 0.24<sup>a</sup> |
| **Induction** | 202.3 ± 5.39<sup>b</sup> | 180.1 ± 4.38<sup>c</sup> | 20.17 ± 2.93<sup>c</sup> | 8.97 ± 1.37<sup>c</sup> | 6.1 ± 0.39<sup>c</sup> |
| **Induction treated with BER-chloride** | 200.5 ± 8.02<sup>b</sup> | 97.7 ± 5.61<sup>b</sup> | 15.67 ± 2.42<sup>b</sup> | 3.76 ± 0.48<sup>b</sup> | 5.1 ± 0.18<sup>b</sup> |

Values represent the mean ± SD of six rats. ANOVA (one way) followed by Student-Newman-keuls test.
Means with letters (a), (b) and (c) were statistical represented compared to sham control group as follow: a= p < 0.001, b= p < 0.01, c= p < 0.05.

#### 3.1.2. Lipid profiles parameters

Lipid profile in this study showed significant increase in HFD group than that of sham control where TC increased 1.2-fold and TG, LDL-c and vLDL-c were 0.9-fold increase, while HDL-c was significantly decreased by 43%. The disturbance which occurred in lipid profiles was partially repaired after using BER-chloride as treatment for two weeks (Table 3).

**Table 3.** Effect of BER-chloride on lipid profiles after the treatment of diabetic induced experimental animals.

|  | TC (mg/dl) | TG (mg/dl) | HDL-c (mg/dl) | LDL-c (mg/dl) | vLDL-c (mg/dl) |
| --- | --- | --- | --- | --- | --- |
| **Sham control** | 130.5 ± 2.79<sup>a</sup> | 110.3 ± 3.6<sup>a</sup> | 59.33 ± 3.14<sup>a</sup> | 110.33 ± 3.62<sup>a</sup> | 22.1 ± 0.7<sup>a</sup> |
| **Control vehicle** | 127.3 ± 4.41<sup>a</sup> | 110.0 ± 4.47<sup>a</sup> | 64.3 ± 3.01<sup>a</sup> | 41.0 ± 3.74<sup>a</sup> | 22.3 ± 1.2<sup>a</sup> |
| **Control BER-chloride** | 129.3 ± 2.25<sup>a</sup> | 111.2 ± 2.40<sup>a</sup> | 63.2 ± 2.56<sup>a</sup> | 43.9 ± 4.24<sup>a</sup> | 21.77 ± 0.5<sup>a</sup> |
| **Induction** | 282.2 ± 3.31<sup>c</sup> | 210.5 ± 3.73<sup>c</sup> | 33.3 ± 2.88<sup>c</sup> | 206.7 ± 2.78<sup>c</sup> | 42.9 ± 0.4<sup>c</sup> |
| **Induction treated with BER-chloride** | 164.5 ± 4.14<sup>b</sup> | 141.5 ± 2.26<sup>b</sup> | 42.8 ± 3.37<sup>b</sup> | 93.4 ± 6.20<sup>b</sup> | 28.133 ± 0.7<sup>b</sup> |

Values represent the mean ± SD of six rats. ANOVA (one way) followed by Student-Newman-keuls test.
Means with letters (a), (b) and (c) were statistical represented compared to sham control group as follow: a= p < 0.001, b= p < 0.01, c= p < 0.05.

#### 3.1.3. Oxidative stress markers, serum transaminases activity and kidney function tests

Experimental HFD rats showed elevation of TBARS and XO (1.5 and 0.8-folds) compared to sham control. Both TBARS and XO were decreased nearly 0.4 and 0.2 folds after two weeks from BER-chloride treatment. On the other hand, the GSH, GPx and ATPase were decreased by 50.9%, 41.9% and 35.2% in HFD group compared with sham control and after administration of BER-chloride; those previous parameters were increased by percentage 42.9, 33.7 and 26.1 respectively (Table 4). Also, HFD intake increased both liver function parameters and kidney function test comparing to sham control one. Administration of 100 mg/kg Bwt BER-chloride in HFD rats for two weeks leads to reduction of liver enzymes activities and kidney function tests nearly 0.3 and 0.2-folds (Table 5).

**Table 4.**
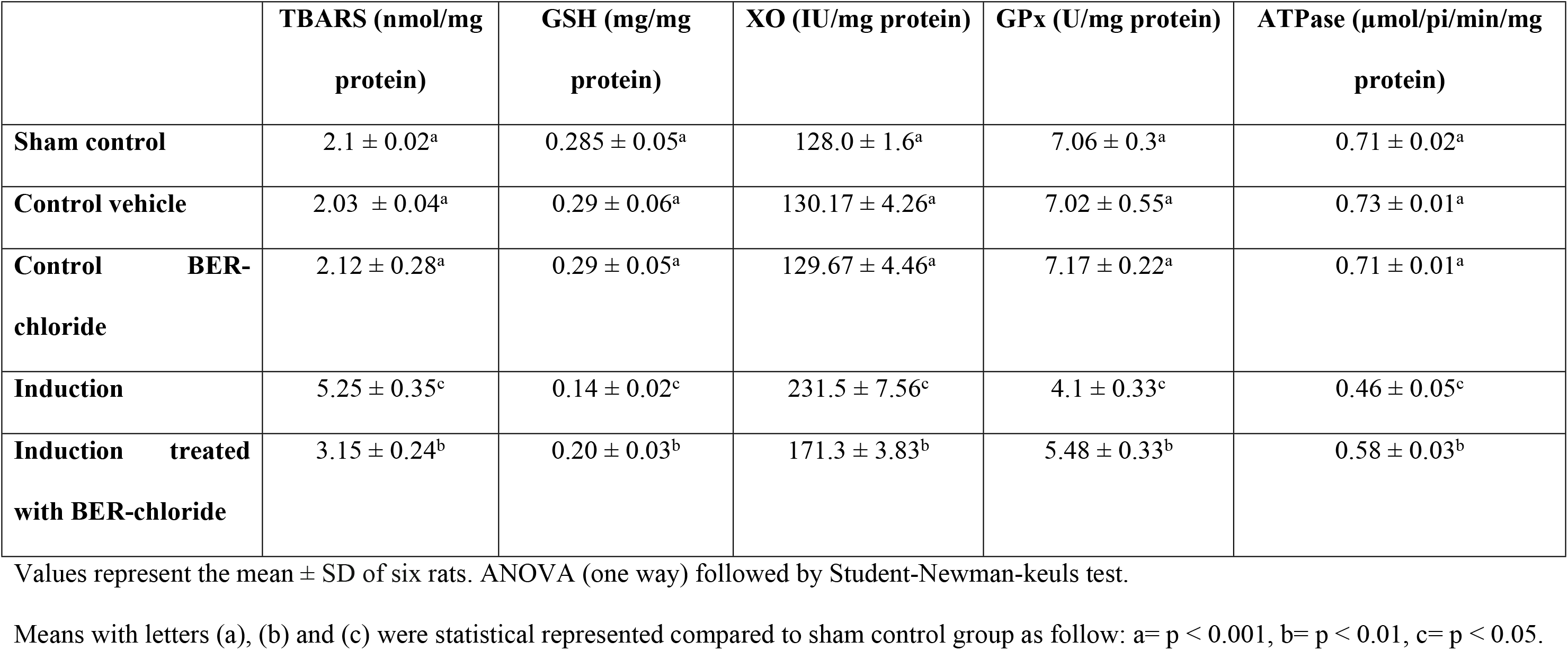
Effect of BER-chloride on hepatocyte prooxidants/antioxidants status after the treatment of diabetic induced experimental animals.

|  | **TBARS (nmol/mg**<br>**protein)** | **GSH (mg/mg**<br>**protein)** | **XO (IU/mg protein)** | **GPx (U/mg protein)** | **ATPase (μmol/pi/min/mg**<br>**protein)** |
| --- | --- | --- | --- | --- | --- |
| **Sham control** | 2.1 ± 0.02<sup>a</sup> | 0.285 ± 0.05<sup>a</sup> | 128.0 ± 1.6<sup>a</sup> | 7.06 ± 0.3<sup>a</sup> | 0.71 ± 0.02<sup>a</sup> |
| **Control vehicle** | 2.03 ± 0.04<sup>a</sup> | 0.29 ± 0.06<sup>a</sup> | 130.17 ± 4.26<sup>a</sup> | 7.02 ± 0.55<sup>a</sup> | 0.73 ± 0.01<sup>a</sup> |
| **Control BER-**<br>**chloride** | 2.12 ± 0.28<sup>a</sup> | 0.29 ± 0.05<sup>a</sup> | 129.67 ± 4.46<sup>a</sup> | 7.17 ± 0.22<sup>a</sup> | 0.71 ± 0.01<sup>a</sup> |
| **Induction** | 5.25 ± 0.35<sup>c</sup> | 0.14 ± 0.02<sup>c</sup> | 231.5 ± 7.56<sup>c</sup> | 4.1 ± 0.33<sup>c</sup> | 0.46 ± 0.05<sup>c</sup> |
| **Induction treated**<br>**with BER-chloride** | 3.15 ± 0.24<sup>b</sup> | 0.20 ± 0.03<sup>b</sup> | 171.3 ± 3.83<sup>b</sup> | 5.48 ± 0.33<sup>b</sup> | 0.58 ± 0.03<sup>b</sup> |

Values represent the mean ± SD of six rats. ANOVA (one way) followed by Student-Newman-keuls test.
Means with letters (a), (b) and (c) were statistical represented compared to sham control group as follow: a= p < 0.001, b= p < 0.01, c= p < 0.05.

**Table 5.** Effect of BER-chloride on serum transaminases activities and kidney function tests after the treatment of diabetic induced experimental animals.

|  | **ALT (U/ml)** | **AST (U/ml)** | **Urea (mg/dl)** | **Creatinine (mg/dl)** |
| --- | --- | --- | --- | --- |
| **Sham control** | 33.5 ± 2.1<sup>a</sup> | 46.0 ± 3.2<sup>a</sup> | 33.4 ± 2.3<sup>a</sup> | 0.71 ± 0.1<sup>a</sup> |
| **Control vehicle** | 34.2 ± 4.6<sup>a</sup> | 49.3 ± 1.8<sup>a</sup> | 31.83 ± 4.6<sup>a</sup> | 0.72 ± 0.04<sup>a</sup> |
| **Control BER-chloride** | 32.2 ± 1.94<sup>a</sup> | 49.5 ± 3.78<sup>a</sup> | 29.67 ± 2.16<sup>a</sup> | 0.71 ± 0.04<sup>a</sup> |
| **Induction** | 58.5 ± 5.09<sup>c</sup> | 85.7 ± 5.68<sup>c</sup> | 56.17 ± 2.14<sup>c</sup> | 0.90 ± 0.05<sup>b</sup> |
| **Induction treated with**<br>**BER-chloride** | 40.8 ± 3.71<sup>b</sup> | 62.5 ± 6.80<sup>b</sup> | 39.83 ± 3.49<sup>b</sup> | 0.75 ± 0.06<sup>a</sup> |

Values represent the mean ± SD of six rats. ANOVA (one way) followed by Student-Newman-keuls test.
Means with letters (a), (b) and (c) were statistical represented compared to sham control group as follow: a= p < 0.001, b= p < 0.01, c= p < 0.05.

### 3.2. Insulin signaling pathway parameters and adiponectin concentration

#### 3.2.1. Insulin signaling pathway parameters

Marked significant reduction of PI3K, AKt-p and SIRT-1 by 57.1%, 42.9% and 61.9%, was reported in HFD rats compared to sham control. The treatment of rats with BER-chloride leads to increase PI3K, AKt-p and SIRT-1 to 84.4%, 49% and 120%, respectively. However, PTEN was increased in HFD rats by 66.6% and decreased to 27.5% after BER-chloride treatment (Table 6). Also, RBP4 was decreased from 1.5-fold to 0.4-fold after BER-chloride treatment. However, BER-chloride failed to affect the HFD adverse effect on IR expression (Fig.1).

**Table 6.** Effect of BER-chloride on PI3K, Akt-p, SIRT-1 and PTEN levels in hepatocyte after the treatment of diabetic induced experimental animals.

|  | **PI3K (pg/g)** | **Akt-p (pg/g)** | **SIRT-1 (ng/g)** | **PTEN (pg/g)** |
| --- | --- | --- | --- | --- |
| **Sham control** | 493.0 ± 12.3<sup>c</sup> | 350.0 ± 23.5<sup>c</sup> | 105.0 ± 10.2<sup>c</sup> | 1200±28.9<sup>a</sup> |
| **Control vehicle** | 486.0 ± 12.0<sup>c</sup> | 335.0 ± 19.3<sup>c</sup> | 110.0 ± 6.9<sup>c</sup> | 1250.0 ± 25.3<sup>a</sup> |
| **Control BER-chloride** | 491.0 ± 23.5<sup>c</sup> | 342.0 ± 18.9<sup>c</sup> | 109.0 ± 9.7<sup>c</sup> | 1150.0 ± 65.3<sup>a</sup> |
| **Induction** | 211.5 ± 21.5<sup>a</sup> | 200.0 ± 12.1<sup>a</sup> | 40.0 ± 1.2<sup>a</sup> | 2000.0 ± 28.9<sup>c</sup> |
| **Induction treated with**<br>**BER-chloride** | 390.0 ± 31.2<sup>b</sup> | 298.0 ± 14.2<sup>b</sup> | 86.0 ± 8.5<sup>b</sup> | 1450.0 ± 23.5<sup>b</sup> |

Values represent the mean ± SD of six rats. ANOVA (one way) followed by Student-Newman-keuls test.
Means with letters (a), (b) and (c) were statistical represented compared to sham control group as follow: a= p < 0.001, b= p < 0.01, c= p < 0.05.

#### 3.2.2. Adiponectin concentration

Adiponectin percentage in serum, liver and white adipose tissue of HFD rats was reduced (52%, 80% and 45%), and this percentage was elevated after two weeks of BER-chloride treatment (48%, 385% and 65.3%), respectively (Table 7).

**Table 7.** Effect of BER-chloride on adiponectin level in serum, liver and white adipose tissue homogenates after the treatment of diabetic induced experimental animals.

|  | Adiponectin (ng/ml) |  |  |
| --- | --- | --- | --- |
|  | Serum | Liver | White adipose tissue |
| **Sham control** | 1.42 ± 0.49<sup>a</sup> | 11.03 ± 0.97<sup>a</sup> | 21.42 ± 1.65<sup>a</sup> |
| **Control vehicle** | 1.39 ± 0.49<sup>a</sup> | 10.67 ± 0.97<sup>a</sup> | 22.97 ± 1.12<sup>a</sup> |
| **Control BER-chloride** | 1.59 ± 0.26<sup>a</sup> | 10.77 ± 1.15<sup>a</sup> | 23.43 ± 1.51<sup>a</sup> |
| **Induction** | 0.682 ± 0.11<sup>b</sup> | 2.0 ± 0.7<sup>b</sup> | 11.78 ± 1.89<sup>b</sup> |
| **Induction treated with BER-chloride** | 1.01 ± 0.18<sup>a</sup> | 9.7 ± 0.9<sup>a</sup> | 19.47 ± 1.67<sup>a</sup> |

Values represent the mean ± SD of six rats. SPSS (Version 20.0).
Means with letters (a), (b) and (c) were statistical represented compared to sham control group as follow: a= p < 0.001, b= p < 0.01, c= p < 0.05.

### 3.3. Histological results

The biochemical results were confirmed by the histological studies in white adipose tissue (Figs. 2A-2E). Control rat’s white adipose tissue revealed normal tissue (Fig 2A). Both PEG and BER-chloride administrated groups after feeding LFD were similar to control rats (Fig 2B and 2C). However, adipose tissue of HFD rats revealed multiple fibrosis and degeneration for the architecture of the adipocytes (Fig 2D). Treatment of HFD rats with BER-chloride for two weeks lead to regeneration of the cells and reduction of the lipids droplets inside it (Fig 2E).

**Fig 2.**
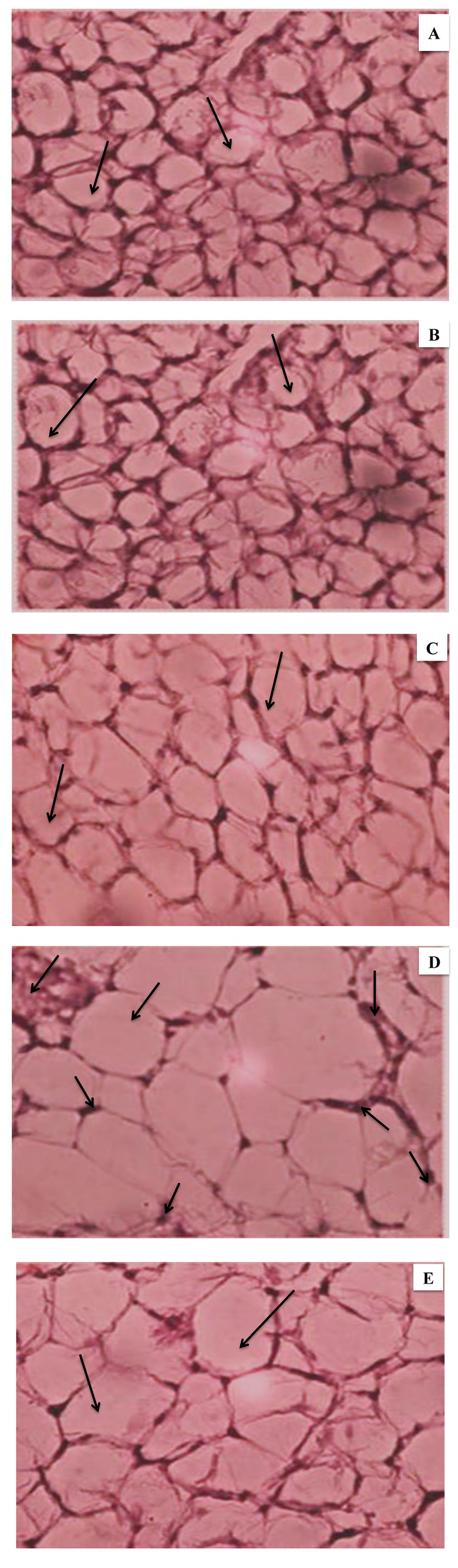
White adipocyte sections pictures in the different groups of rats, stained with hematoxelin and eosin. (A) Sham control rats, (B) Control vehicle (PEG) rats, (C) Control BER-chloride, (D) HFD-fed rats and (E) HFD-fed rats and treated with BER-chloride (X = 400).

## 4. Discussion

In recent years insulin resistance has a huge challenge, it represented one of the risk metabolic conditions, and in many cases it occurs due to bad dietary habits such as junk foods which is characterized by huge percentage of fats [24] or due to some diseases like non-insulin dependent diabetes mellitus [25]. In present study HFD feeding for eight weeks induced hyperinsulinaemia which is attributed to inability of liver to utilize the secreted insulin although the normal function of pancreatic β-cell [26]. During this period the rats become more obese compared to LFD controls due to elevation of insulin level which inhibit fatty acid oxidation so fats accumulated mainly in liver because it is the main organ of oxidation process [27]. Elevated levels of fasting insulin, glucose, and HOMA-IR index confirming the state of insulin resistance. Furthermore increasing of HbA1c is another indicator of insulin resistance and correlated with renal function parameters elevation [28].

Additionally, results of current study showed up regulation of RBP4 expression in HFD rats. From previous studies RBP4 i: elevates the process of gluconeogenesis in liver so hyperglycemia is occurred [29], ii: hindered the insulin signaling in muscle and iii: decreased the uptake of glucose by reduction the activity of PI3K [30]. Defects in PI3K demonstrated in other findings [31]. Previous studies showed highest level of RBP4 are associated with body mass index increase, insulin resistance and hypertriglyceridemia [32].

Study of PTEN in this research had a significant importance. PTEN is lipid phosphatase inhibits insulin signaling by dephosphorylating phosphatidylinositol (3,4,5) triphosphate (PIP_3_) to phosphatidylinositol (4,5) diphosphate (PIP_2_) [33]. Hence, PTEN is antagonizing the action of PI3K and inhibits Akt as appear in current results, where HFD rats showed elevation in PTEN level and reduction of PI3K [34]. It is known that, SIRT1 is a key regulator of lipid mobilization through its action together with adenosine monophosphate (AMPK) by increasing fatty acid metabolism [35]. So, reduction of SIRT1 concentration in HFD rats is linked by hyperlipidemia and insulin resistance due to decrease in phosphorylated and/or activated AMPK resulting in lipid synthesis increase [36]. Results of current study are in accordance with previous studies [37]. Further, SIRT1 has several roles in insulin signaling pathway it i: regulates secretion of insulin from β-cell of pancreas by reduction the expression of uncoupling protein-2 (UCP2) and improvement the depolarization in β-cell of pancreas [38] and ii: regulates the insulin signaling pathway by deacetylation of insulin receptor substrate-2 (IRS2) and activation of Akt in cells [39]. From those mentioned mechanisms of SIRT1 and PTEN, the current study showed reduction of Akt-p concentration in hepatic cells of HFD rats.

Moreover adiponectin has an important role in insulin resistance pathway; it is a member of adipocytokines which secreted by adipocytes and has a regulating effect on insulin sensitivity [40]. It was reported that disturbance in lipid metabolism and excessive fat deposition leads to abnormal synthesis of adipocytokines [41]. HFD rats associated with reduction of adiponectin level in serum, hepatic and adipose tissues which may be attributed to disruption of both adiponectin receptor-1 and 2 leading to elevation of glucose level and reduction the activity of peroxisome proliferator activated receptor α-signaling pathways respectively, and finally insulin resistance occurs [42]. Further, there are some studies suggest the role of SIRT1 in regulation of adiponectin secretion from the adipocytes by deacetylating of fork head transcription factor O1 (FOXO-1) protein and enhancement the transcription of gene that encodes adiponectin in adipocytes [43]. Hence, reduction of SIRT1 effect on adiponectin secretion.

Role of SIRT1 is extended to control the production of reactive oxygen species (ROS) [44] as SIRT1 is considered one of the important proteins that protect cells from stress damage [37]. Under normal condition the hepatocyte balance the oxidative stress by the action of antioxidant enzymes such as GPx which converts hydrogen peroxide (H_2_O_2_) to water [45]. Rats suffer from insulin resistance have low GPx activity so H_2_O_2_ accumulated and hepatic cells damaged. These results were confirmed by elevation of liver enzymes (ALT and AST). H_2_O_2_ accumulation also affected on renal tissue which confirmed by increase both urea and creatinine levels. Another cause of elevation of ROS in case of insulin resistance is attributed to the dysregulated production of adipocytokines where plasma adiponectin concentration is inversely correlated with systemic oxidative stress [46].

In recent years with regard to the adverse effects of synthetic drugs, increasing attention has been paid by researchers to herbal medicines. BER is a major form of isoquinoline alkaloid isolated from several herbal plants and it has several biological effects [47]. Nowadays, BER is manufactured by chemical synthesis, chloride or sulfate salt of BER is used for clinical purposes [48].

Our *in vivo* study revealed that treatment with BER-chloride has negative effect on insulin resistance by activating two proteins involved in several physiological processes, SIRT-1 and AMPK [49]. Those two proteins able to activate each other, AMPK activates SIRT-1 by elevation the level of nicotinamide phosphoribosyltransferase and SIRT-1 stimulates AMPK through deacetylation of serine-thereonine kinase LKB1 [50, 51]. Also, BER has the ability to improve insulin resistance through other mechanisms, where it i: protects β-cell of islet of *Langerhans* from damage, ii: allows glucose uptake of skeletal muscle, iii: improves hepatic gluconeogenesis and iv: decreases the level of lipids in blood [52, 53].

As a result of SIRT-1 and AMPK pathways activation, adiponectin level was restored after BER-chloride administration in our study which similar to results obtained by [54]. Elevation of adiponectin level is linked by regulation of β-oxidation of fatty acids and glucose metabolism [55, 56]. Hence, treated rats with BER-chloride showed significant reduction of lipid profiles and BGL. Also, BER-chloride ameliorates hyperlipidemia results from insulin resistance via different mechanisms i: BER lowers blood cholesterol levels through inhibiting cholesterol uptake and absorption in the intestine [57], ii: BER reduces the secretion of cholesterol from enterocytes into the blood by down regulation acetyl CoA transferase II enzyme [58] and lastly iii: BER increases the regulation of LDL-receptor and hence, BER decreases the level of LDL-c [59]. Further, BER chloride is able to improve glucose control by stimulation the glycolysis in peripheral tissue [60], inhibition of FOXO-1 and hepatic nuclear factor 4, lead to suppression of glucose-6-phosphatase and phosphoenolpyruvate carboxykinase enzymes which responsible for liver gluconeogenesis [61, 62] and activation of glucose transport-1 (GLUT1) [63].

The current study showed BER-chloride has the ability to increase the level of antioxidant enzymes (GSH, GPx and XO) by reducing the elevated level of lipid peroxidation [64]. Also it was reported that BER has the ability to prevent nicotinamide adenine dinucleotide phosphate (NADPH) oxidase which is a major source of ROS production [65]. These results in accordance with previous studies which proved that BER is a strong antioxidant molecule due to its ability to scavenge free radicals [66]. Moreover, BER-chloride exerts protective effect against ROS through SIRT-1 activation where SIRT-1 able to modulate NOX4/NADPH oxidative subunit [67]. Reduction of ROS production lead to decrease the level of liver enzymes in group of rats administrated BER-chloride.

In current study, BER-chloride down regulates RBP4 which acts as an effective insulin sensitizing function [68]. From previous studies, it was reported that reduction the level of RBP4 is related to elevation of HDL-c and decrease TG levels in some patients [69]. Also as result of reduction of SIRT1 in HFD rats, elevation of TBARS and XO and reduction of GPx and GSH was noticed and this was improved after BER-chloride administration.

## 5. Conclusions

Berberine chloride can be considered one of therapeutics used to decrease insulin resistance through its effect on several insulin signaling pathways. A schematic representation was designed (Fig 3) to summarize the modification of insulin resistance by BER-chloride.

**Fig 3.**
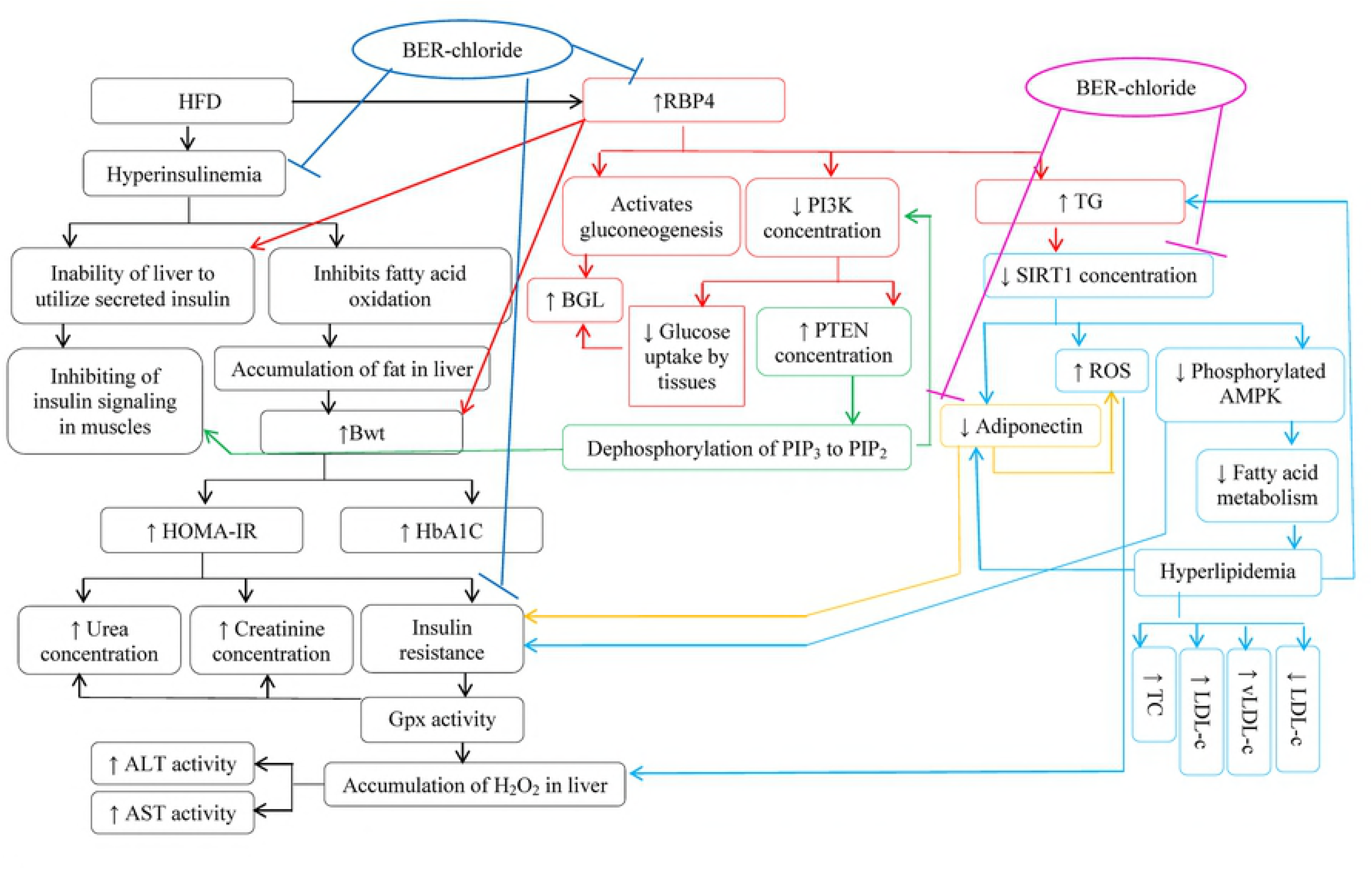
Schematic diagram for the effect of BER-chloride on insulin signaling in HFD-insulin resistance induced rats. Blue arrows 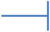 means inhibition and violet arrows 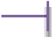 means activation.

## Acknowledgements

This work is financially supported by Alexandria University, Egypt.

## References

1. Steven EK, Rebecca LH, Kristina MU. Mechanisms linking obesity to insulin resistance and type 2 diabetes. Nature. 2006;444: 840-846.

2. Joshua AD, William JR, Piul SR, Daniel JC. The Nrf2/Keap1/ARE pathway and oxidative stress as a therapeutic target in type II diabetes mellitus. J Diabetes Res. 2017;2017: 1-15.

3. Gao YF, Zhang MN, Wang TX, Wu TC, Ai RD, Zhang ZS. Hypoglycemic effect of D-chiro-inositol in type 2 diabetes mellitus rats through the PI3K/Akt signaling pathway. Mol Cell Endocrinol. 2016;433: 26-34.

4. Jiang WX, Li SH, Li XJ. Therapeutic potential of berberine against neurodegenerative diseases. Life Sci. 2015;58: 564-569.

5. Bodiwala HS, Sabde S, Mitra D, Bhutani KK. Synthesis of 9-substituted derivatives of berberine as anti-HIV agents. Eur J Med Chem. 2011;46: 1045-1049.

6. Li J-W, Yuan K, Shang S-C, Guo Y. A safer hypoglycemic agent for type 2 diabetes-Berberine organic acid salt. J Funct Foods. 2017;38: 399-408.

7. Liu Q, Xu X, Zhao M, Wei Z, Li X, Zhang X, et al. Berberine induces senescence of human glioblastoma cells by down regulating the EGFR-MEK-ERK signaling pathway. Mol Cancer Ther. 2015;14: 355-363.

8. Vuddanda PR, Chakraborty S, Singh S. Berberine: A potential phytochemical with multispectrum therapeutic activities. Expert Opin Investig Drugs. 2010;19: 1297-1307.

9. Zhang S, Wang X, Yin W, Liu Z, Zhou M, Xiao D, Liu Y, Peng, D. Synthesis and hypoglycemic activity of 9-O-(lipophilic group substituted) berberine derivatives. Bioorg Med Chem Lett. 2016;26: 4799-4803.

10. Dong SF, Hong Y, Liu M, Hao YZ, Yu HS, Liu Y, Sun JN. Berberine attenuates cardiac dysfunction in hyperglycemic and hypercholesterolemic rats. Eur J Pharmacol. 2011;660: 368-374.

11. El-Sayed M, Ghareeb D, Talat H, Sarhan E. High fat diet induced insulin resistance and elevated retinol binding protein 4 in female rats; treatment and protection with *Berberis vulgaris* extract and vitamin A. PJPS. 2013;26: 1189-1195.

12. Ghareeb D, Khalil S, Hafez H, Bajorath J, Ahmed H, Sarhan E, Elwakeel E, El-Demellawy M. Berberine reduces neurotoxicity related to nonalcoholic steatohepatitis in rats. Evid Based Complement Alternat Med. 2015;2015: 1-13.

13. Saleh SR, Attia R, Ghareeb DA. The ameliorating effect of *Berberine*-rich fraction against gossypol-induced testicular inflammation and oxidative Stress. Oxid Med Cell Longev. 2018;2018. 1056173.

14. Steinberg D, Vaughan M, Margolis S. Studies of triglyceride in homogenates of adipose tissue. Biol Chem. 1961;236: 1631-1637.

15. Anuradha CV, Pavikumar P. Anti-lipid peroxidative activity of seeds of fenugreek *(Trigonella foenum graecum)*. Med Sci Res. 1998;26: 317-321.

16. Han SJ, Boyko EJ, Kim S-K, Fujimoto WY, Kahn, SE, Leonetti, DL. Association of thigh muscle mass with insulin resistance and incident type 2 diabetes mellitus in Japanese Americans. Diabetes Metab J. 2018;42: 1-8.

17. Friedewald WT, Levy RI, Fredrickson DS. Estimation of the concentration of low density lipoprotein cholesterol in plasma without use of the preparative ultracentrifuge. Clin Chem. 1972; 18: 499-502.

18. Tappel AL, Zalkin H. Inhibition of lipid peroxidation in mitochondria by vitamin E. Arch. Biochem. Biophys. 1959;80: 333-336.

19. Ellman GL. Tissue sulfhydryl groups. Arch Biochem Biophys. 1959; 82: 70-77.

20. Litwack G, Bothwell JW, Williams JN, Elvehjem Jr CA. A colorimetric assay for xanthine oxidase in rat liver homogenates. J Biol Chem. 1953;200: 303-310.

21. Paglia E, Valentine N. Studies on the quantitative and qualitative characterization of erythrocyte glutathione peroxidase. J Lab Clin Med. 1967;70: 158-169.

22. Chiu DTY, Stults FH, Tappel AL. Purification and properties of rat lung soluble glutathione peroxidase. Biochim. Biophys. Acta. 1976;445: 558-566.

23. Candeias MF, Abreu P, Pereira A, Cruz-Morais J. Effects of strictosamide on mouse brain and kidney Na^+^, K^+^-ATPase and Mg^2+^-ATPase activities. J Ethnopharmacol. 2009;121: 117-122.

24. Haslam DW, James WP. Obesity. Lancet. 2005;366: 1197-1209.

25. Seko Y, Sumida Y, Tanaka S, Mori K, Taketani H, Ishiba H, et al. Insulin resistance increases the risk of incident type 2 diabetes mellitus in patients with non-alcoholic fatty liver disease. Hepatol Res. 2018;48: E42-E51.

26. Arnold SE, Arvanitakis Z, Macauley-Rambach SL, Koenig AM, Wang H-Y, Ahima RS, et al. Brain insulin resistance in type 2 diabetes and Alzheimer disease: concepts and conundrums. Nat Rev Neurol. 2018;14: 168-181.

27. Liu R, Li H, Fan W, Jin Q, Chao T, Wu Y, et al. Leucine supplementation differently modulates branched-chain amino acid catabolism, mitochondrial function and metabolic profiles at the different stage of insulin resistance in rats on high-fat diet. Nutrients. 2017;9: 565-585.

28. Fiorentino TV, Marini MA, Succurro E, Sciacqua A, Andreozzi F, Perticone F, Sesti G. Elevated hemoglobin glycation index identify non-diabetic individuals at increased risk of kidney dysfunction. Oncotarget. 2017;8: 79576-79586.

29. Hutchison SK, Harrison C, Stepto N, Meyer C, Teede HJ. Retinol binding protein 4 and insulin resistance in polycystic ovary syndrome. Diabetes Care. 2008;31: 1427-1432.

30. Yousefi MR, TaheriChadorneshin H. The effect of moderate endurance training on gastrocnemius retinol-binding protein 4 and insulin resistance in streptozotocin-induced diabetic rats. Interv Med Appl Sci. 2017;9: 1-5.

31. Jiang G, Zhang BB. Pi 3-kinase and its up- and down-stream modulators as potential targets for the treatment of type II diabetes. Front. Biosci. 2002;7: d903 - d907.

32. Domingos MAM, Queiroz M, Lotufo PA, Benseñor IJ, Titan SMO. Serum RBP4 and CKD: Association with insulin resistance and lipids. J Diabetes Complications. 2017;31: 1132-1138.

33. Cully M, You H, Levine AJ, Mak TW. Beyond PTEN mutations: the PI3K pathway as an integrator of multiple inputs during tumorigenesis. Nat Rev Cancer. 2006;6: 184-192.

34. Yao XH, Nyomba BLG. Hepatic insulin resistance induced by prenatal alcohol exposure is associated with reduced PTEN and TRB3 acetylation in adult rat offspring. Am J Physiol Regul Integr Comp Physiol. 2008;294: R1797-R1806.

35. Merksamer PI, Liu Y, He W, Hirschey MD, Chen D, Verdin E. The sirtuins, oxidative stress and aging: an emerging link. Aging (Albany NY). 2013;5: 144-150.

36. Boulet MM, Chevrier G, Grenier-Larouche T, Pelletier M, Nadeau M, Scarpa J. et al. Alterations of plasma metabolite profiles related to adipose tissue distribution and cardiometabolic risk. Am J Physiol Endocrinol Metab. 2015;309: E736-E746.

37. Deng XQ, Chen LL, Li NX. The expression of SIRT1 in nonalcoholic fatty liver disease induced by high-fat diet in rats. Liver Int. 2007;27: 708-715.

38. Liang F, Kume S, Koya D. SirT1 and insulin resistance. Nat Rev Endocrinol. 2009;5: 367-373.

39. Yoshizaki T, Milne JC, Imamura T, Schenk S, Sonoda N, Babendure JL. SirT1 exerts anti-inflammatory effects and improves insulin sensitivity in adipocytes. Mol. Cell Biol. 2009;29: 1363-1374.

40. Wu Q-M, Ni H-X, Lu X. Changes of adipocytokine expression after diabetic rats received sitagliptin and the molecular mechanism. Asian Pac J Trop Med. 2016;9: 893-897.

41. Boulet MM, Chevrier G, Grenier-Larouche T, Pelletier M, Nadeau M, Scarpa J. et al. Alterations of plasma metabolite profiles related to adipose tissue distribution and cardiometabolic risk. Am J Physiol Endocrinol Metab. 2015;309: E736-E746.

42. Li S, Shin HJ, Ding EL, van Dam RM. Adiponectin levels and risk of type 2 diabetes. A systematic review and meta-analysis. JAMA. 2009;302: 179-188.

43. Qiao L, Shao J. SirT1 regulates adiponectin gene expression through Foxo1-C/enhancer binding protein alpha transcriptional complex. J Biol Chem. 2006;281: 39915-39924.

44. Colak Y, Ozturk O, Senates E, Tuncer I, Yorulmaz E, Adali G, et al. SIRT1 as a potential therapeutic target for treatment of nonalcoholic fatty liver disease. Med Sci Monit. 2011;17: HY5-HY9.

45. Flores C, Adhami N, Martins-Green M. THS toxins induce hepatic steatosis by altering oxidative stress and SIRT1 levels. J Clin Toxicol. 2016;6: 318.

46. Furukawa S, Fujita T, Shimabukuro M, Iwaki M, Yamada Y, Nakajima Y, et al. Increased oxidative stress in obesity and its impact on metabolic syndrome. J Clin Invest. 2004;114: 1752-1761.

47. Jin Y, Khadka DB, Cho WJ. Pharmacological effects of berberine and its derivatives: a patent update. Expert Opin Ther Pat. 2016;26: 229 - 243.

48. Battu SK, Repka MA, Maddineni S, Chittiboyina AG, Avery MA. Majumdar S. Physicochemical characterization of berberine chloride: aperspective in the development of a solution dosage form for oral delivery. AAPS PharmSciTech. 2010;11: 1466-1475

49. Lin Y, Sheng M, Ding Y, Zhang N, Song Y, · Du H, et al. Berberine protects renal tubular cells against hypoxia / reoxygenation injury via the Sirt1/p53 pathway. J Nat Med. 2018;72: 715-723.

50. Sun Y, Li J, Xiao N, Wang M, Kou J, Qi L, et al. Pharmacological activation of AMPK ameliorates perivascular adipose/endothelial dysfunction in a manner interdependent on AMPK and SIRT1. Pharmacol Res. 2014;89: 19-28.

51. Hardie DG. AMPK: positive and negative regulation, and its role in whole-body energy homeostasis. Curr Opin Cell Biol. 2015;33: 1-7.

52. Yang TC, Chao HF, Shi LS, Chang TC, Lin HC, Chang WL. Alkaloids from *Coptis chinensis* root promote glucose uptake in C2C12 myotubes. Fitoterapia. 2014;93: 239-244.

53. Pirillo A, Catapano AL. Berberine, a plant alkaloid with lipid-and glucose-lowering properties: From *in vitro* evidence to clinical studies. Atherosclerosis. 2015;243: 449-461.

54. Wu Y, Cha Y, Huang X, Liu J, Chen Z, Wang F, et al. Protective effects of berberine on high fat-induced kidney damage by increasing serum adiponectin and promoting insulin sensitivity. Int J Clin Exp Pathol. 2015;8: 14486-14492.

55. Yamauchi T, Kamon J, Minokoshi Y, Ito Y, Waki H, Uchida S, et al. Adiponectin stimulates glucose utilization and fatty-acid oxidation by activating AMP activated protein kinase. Nat. Med. 2002;8: 1288-1295.

56. Lin H, Li Z. Adiponectin self-regulates its expression and multimerization in adipose tissue: An autocrine/paracrine mechanism? Med Hypotheses. 2012;78: 75-78.

57. Wang Y, Yi X, Ghanam K, Zhang S, Zhao T, Zhu X. Berberine decreases cholesterol levels in rats through multiple mechanisms, including inhibition of cholesterol absorption. Metb Clin Exp. 2014;63: 1167-1177.

58. Wang H, Zhu C, Ying Y, Luo L, Huang D, Luo Z. Metformin and berberine, two versatile drugs in treatment of common metabolic diseases. Oncotarget. 2018;9: 10135-10146.

59. Abidi P, Zhou Y, Jiang JD, Liu J. Extracellular signal-regulated kinase-dependent stabilization of hepatic low-density lipoprotein receptor mRNA by herbal medicine berberine. Arterioscler Thromb Vasc Biol. 2005;25: 2170-2176.

60. Yin J, Gao Z, Liu D, Liu Z, Ye J. Berberine improves glucose metabolism through induction of glycolysis. Am J Physiol Endocrinol Metab. 2008;294: 148-156.

61. Kim WS, Lee YS, Cha SH, Jeong HW, Choe SS, Lee MR, et al. Berberine improves lipid dysregulation in obesity by controlling central and peripheral AMPK activity. Am J Physiol Endocrinol Metab. 2009;296: 812-819.

62. Xia X, Yan J, Shen Y, Tang K, Yin J, Zhang Y, et al. Berberine improves glucose metabolism in diabetic rats by inhibition of hepatic gluconeogenesis. PLoS. One. 2011; 6: 16556.

63. Cok A, Plaisier C, Salie MJ, Oram DS, Chenge J, Louters LL. Berberine acutely activates the glucose transport activity of GLUT1. Biochimie. 2011;93: 1187-1192.

64. Lao-ong T, Chatuphonprasert W, Nemoto N, Jarukamjorn K. Alteration of hepatic glutathione peroxidase and superoxide dismutase expression in streptozotocin-induced diabetic mice by berberine. Pharm Biol. 2012;50: 1007-1012.

65. Cheng F, Wang Y, Li J, Su C, Wu F, Xia WH. et al. Berberine improves endothelial function by reducing endothelial microparticles-mediated oxidative stress in humans. Int J Cardiol. 2013;167: 936-942.

66. Kumar A, Ekavali, Chopra K, Mukherjee M, Pottabathini R, Dhull DK. Current knowledge and pharmacological profile of berberine: An update. Eur J Pharmacol. 2015;761: 288-297.

67. Karbasforooshan H, Karimi G. The role of SIRT1 in diabetic retinopathy. Biomed. Pharmacother. 2018;97: 190-194.

68. Zhang W, Xu YC, Guo FJ, Meng Y, Li ML. Antidiabetic effects of cinnamaldehyde and berberine and their impacts on retinol-binding protein 4 expression in rats with type 2 diabetes mellitus. Chin Med J. 2008;121: 2124-2128.

69. Broch M, Gomez JM, Auguet MT, Vilarrasa N, Pastor R, Elio I, et al. Association of retinol-binding protein-4 (RBP4) with lipid parameters in obese women. Obes Surg. 2010;20: 1258-1264.

